# Excitable dynamics in a molecularly-explicit model of cell motility: Mixed-mode oscillations and beyond

**DOI:** 10.1101/2022.10.28.514275

**Authors:** Lucie Plazen, Anmar Khadra

**Affiliations:** Department of Mathematics and Statistics, McGill University, Montreal, Canada; Department of Physiology, McGill University, Montreal, Canada

**Keywords:** Rho family of GTPases, cellular polarity, mathematical model, hysteresis and bistability, wave-pinning, mixed-mode oscillations, relaxation oscillations, slow-fast analysis, cellular potts model, migration patterns.

## Abstract

Mesenchymal cell motility is mainly regulated by two members of the Rho-family of GTPases, called Rac and Rho. The mutual inhibition exerted by these two proteins on each other’s activation and the promotion of Rac activation by an adaptor protein called paxillin have been implicated in driving cellular polarization comprised of front (high active Rac) and back (high active Rho) during cell migration. Mathematical modeling of this regulatory network has previously shown that bistability is responsible for generating a spatiotemporal pattern underscoring cellular polarity called wave-pinning when diffusion is included. We previously developed a 6D reaction-diffusion model of this network to decipher the role of Rac, Rho and paxillin (along with other auxiliary proteins) in generating wave-pinning. In this study, we simplify this model through a series of steps into an excitable 3D ODE model comprised of one fast variable (the scaled concentration of active Rac), one slow variable (the maximum paxillin phosphorylation rate – turned into a variable) and a very slow variable (a recovery rate – also turned into a variable). We then explore, through slow-fast analysis, how excitability is manifested by showing that the model can exhibit relaxation oscillations (ROs) as well as mixed-mode oscillations (MMOs) whose underlying dynamics are consistent with a delayed Hopf bifurcation. By reintroducing diffusion and the scaled concentration of inactive Rac into the model, we obtain a 4D PDE model that generates several unique spatiotemporal patterns that are relevant to cell motility. These patterns are then characterized and their impact on cell motility are explored by employing the cellular potts model (CPM). Our results reveal that wave pinning produces purely very directed motion in CPM, while MMOs allow for meandering and non-motile behaviours to occur. This highlights the role of MMOs as a potential mechanism for mesenchymal cell motility.

## 1 Introduction

Excitability is a very frequently observed phenomenon in many physiological systems including, but not limited to, cellular and molecular biology, neurophysiology [1, 2] and immunology [3]. It typically refers to the ability of a system to exhibit large excursions in their main variables when a small stimulus is applied. Such large excursions could occur singularly, or could occur periodically in a cyclic fashion [2]. It is usually manifested in four different ways, referred to in the literature as type I, II, III and IV excitability [4]. Type I and II are periodic in nature and are generated by homoclinic and Hopf bifurcations, while type III and IV are singular generated by perturbations away from a stable steady state with the former (the latter) depending on differences in time scales (on crossing the stable manifold of a saddle fixed point) to generate the excursion [5–7]. These basic dynamics become significantly more complex when multiple time scaled are involved.

Cell motility, the spontaneous movement of a cell by energy consumption, is an essential process by which many physiological (e.g., embryonic development, wound healing, inflammation) [8, 9] and pathophysiological (e.g., cancer metastasis) [10, 11] systems depend on. It is generated by the asymmetric polarization of migrating cells with a front (or the leading edge) and a back (or the rear edge) along an axis roughly parallel to the path of migration [12]. The molecular networks regulating cell motility are very prominent examples of systems that exhibit diverse types of excitability properties that can range in complexity and impact. Indeed, some of these molecular networks may be able to exhibit multitude of these excitability properties simultaneously when perturbing one parameter [13]. One example of such behaviour was seen in a molecularly-explicit model describing the dynamics of lamellipodium in fish keratocytes [14]. In this latter system, several migration patterns, including waving, traveling wave pulse and smooth motility, were observed [14] and three types of excitabilities were detected in parameter space: type I, III and IV [14, 15].

Mesenchymal cell motility, similar to that seen in Chinese hamster ovary (CHO-K1) cells, is regulated by a network of over 200 proteins (also called adhosome). It is characterized by the formation of protein complexes, called adhesions [16–18], that allow motile cells to anchor on a substrate and to link the extracellular matrix (ECM) to the actin-cytoskeleton network. Although the molecular network governing adhesion dynamics and motility is very complex in these cells, three key proteins have been implicated in regulating both of them. This includes two members of the Rho family of GTPases, namely, Rac1 and RhoA (referred to as Rac and Rho for the remainder of this study) [19–22], as well as an adhesion and adaptor protein called paxillin [23]. This trimolecular regulatory network involves Rac and Rho cycling between their inactive (GDP-bound) to active (GTP-bound) forms, via Guanine nucleotide Exchange Factors (GEFs), and vice versa, via GTPase-Activating Proteins (GAPs) [24], as well as the mutual inhibition that the active forms of these two proteins exert on each other [24]. Active Rac and Rho are membrane bound, making their diffusion coefficients significantly lower than those associated with their inactive forms which happen to be cytosolic. The adhesion protein paxillin, on the other hand, can get phosphorylated at its serine 273 (S273) residue by the active form of the protein p21-Activated Kinase 1 (PAK) when bound to RacGTP (PAK-RacGTP), and subsequently promote Rac activation as a GEF [23,25]. The binding of the protein complex (GIT-PIX-PAK), formed by the G protein-coupled receptor kinase InteracTor 1 (GIT), beta-PAK-Interacting eXchange factor (PIX), and PAK, to phosphorylated paxillin allows this entire complex to act as a GEF [23]. A molecularly explicit reaction-diffusion model of this protein regulatory network was developed in the form of a six-dimensional (6D) partial differential equation (PDE) model [26]. The model described the spatiotemporal dynamics of the scaled concentrations of active/phosphorylated and inactive/unphosphorylated forms of the proteins involved, as well as those auxiliary ones (i.e., GIT, PIX and PAK) that were set to steady states. The model was nondimensionalized, to reduce the number of parameters, and its steady state dynamics were analyzed in the absence of diffusion, demonstrating that it possessed two different time scales. In the presence of diffusion, the model was shown to exhibit wave pinning, a phenomenon in which a travelling wave is pinned in space, generating a concentration gradient in Rac and Rho responsible for forming cellular polarization.

In the absence of diffusion and by taking into account conservation of matter, the model was turned into a 3D ODE model (referred to hereafter as the original ODE model). This original ODE model was shown to possess two timescales and to exhibit a bistable switch. Bistability was defined by the coexistence of two steady states, one corresponding to elevated scaled concentration of active Rac (or low scaled concentration of active Rho) and another corresponding to the low scaled concentration of active Rac (or elevated scaled concentration of active Rho), and was delimited by saddle-node bifurcations. Beyond the saddle-node bifurcations, the model was monostable. The effects of bistability on the dynamics of the 6D PDE was then explored to show that it was responsible for generating wave-pinning with active Rac accumulating in the “front” of the one-dimensional spatial domain, and active Rho accumulating in the back. This phenomenon of wave-pinning produced by the Rac-Rho system was heavily investigated dynamically using a method called local perturbation analysis [27], in which the large difference in diffusion coefficients between the active and inactive forms of Rac and Rho [28] allowed the PDE model to be simplified into an ODE model that can provide insights into how the spatial patterns exhibited by the PDE model after a perturbation is formed. These analyses were helpful in determining the effects of different parameter regimes in defining dynamics.

It is important to point out that other complex models of the Rac-Rho system were also developed by coupling them to the ECM to generate oscillatory dynamics in the absence of diffusion [29]. The amplitude of the oscillations in such models were typically uniform or changed minimally, allowing the PDE models to produce interesting spatiotemporal patterns. Implementing these models using the cellular potts model (CPM), a computer-based discrete grid-based simulation technique that involves the modelling of the ECM as a mesh upon which simulated cells are superimposed [30], typically produced migrating cells that were either purely directional or do not migrate significantly (depending on the amplitude of the oscillations), but not both. These oscillations were consistent with type II excitability through a Hopf bifurcation. In the presence of two competitive ECMs, the oscillations became more “random” (due to the presence of a period doubling bifurcation), but the amplitude of such oscillations did not change significantly change. As a result, the migration patterns produced by this model typically did not succeed in combining both exploratory and stationary behaviours in one simulation typically seen in mesenchymal cell motility [31].

Here, we will expand on these studies by considering series of simplified models of the Rac-Rho-paxillin system capable of producing oscillatory dynamics with nonuniform amplitudes. The models will possess at least two timescales and are excitable in nature, allowing them to exhibit what is known as mixed-mode oscillations (MMOs) [32–34] that combine fast small amplitude oscillations with slow large amplitude oscillations. Because MMOs are more complex dynamically than relaxation oscillations (ROs) and because they are quite frequently observed in models of many physiological systems [32, 35], we will investigate in this study how MMOs impact the spatiotemporal dynamics of the models considered here, especially in comparison to ROs. This will be done numerically by simulating the PDE models whose dynamics are regulated by MMOs and ROs, as well as by employing the CPM to simulate migrating cells.

## 2 The original spatiotemporal 6D PDE model

To understand quantitatively the behaviour of the regulatory network involving Rac, Rho and paxillin, a 6D reaction-diffusion mathematical model describing the dynamics of inactive (GDP bound) and active (GTP bound) forms of Rac and Rho as determined by paxillin S273 phosphorylation was developed [26]. The model consisted of two main modules: the mutual inhibition exerted by Rac and Rho on each other, as well as the indirect positive feedback loop exerted by active Rac on itself through paxillin phosphorylation at S273 (Fig. 1A). The resulting spatiotemporal model over one spatial domain was comprised of 6 PDEs characterizing the interactions of the active and inactive forms of the two proteins Rac (*R, R_i_*) and Rho (*ρ, ρ_i_*) along with the phosphorylated and unphosphorylated forms of paxillin (*P, P_i_*) at S273 residue. Rescaling the model into its nondimensionalized form gave

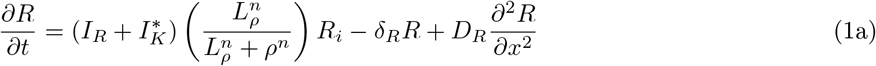

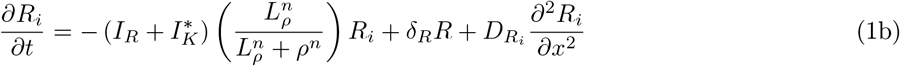

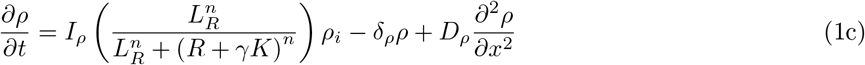

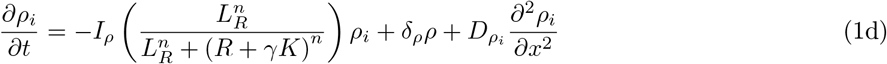

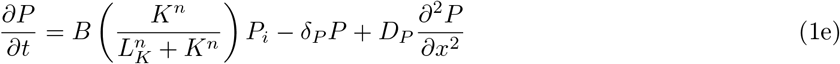

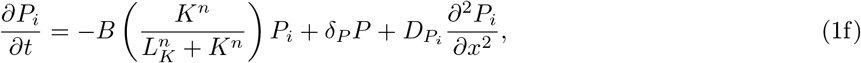

**Figure 1:**
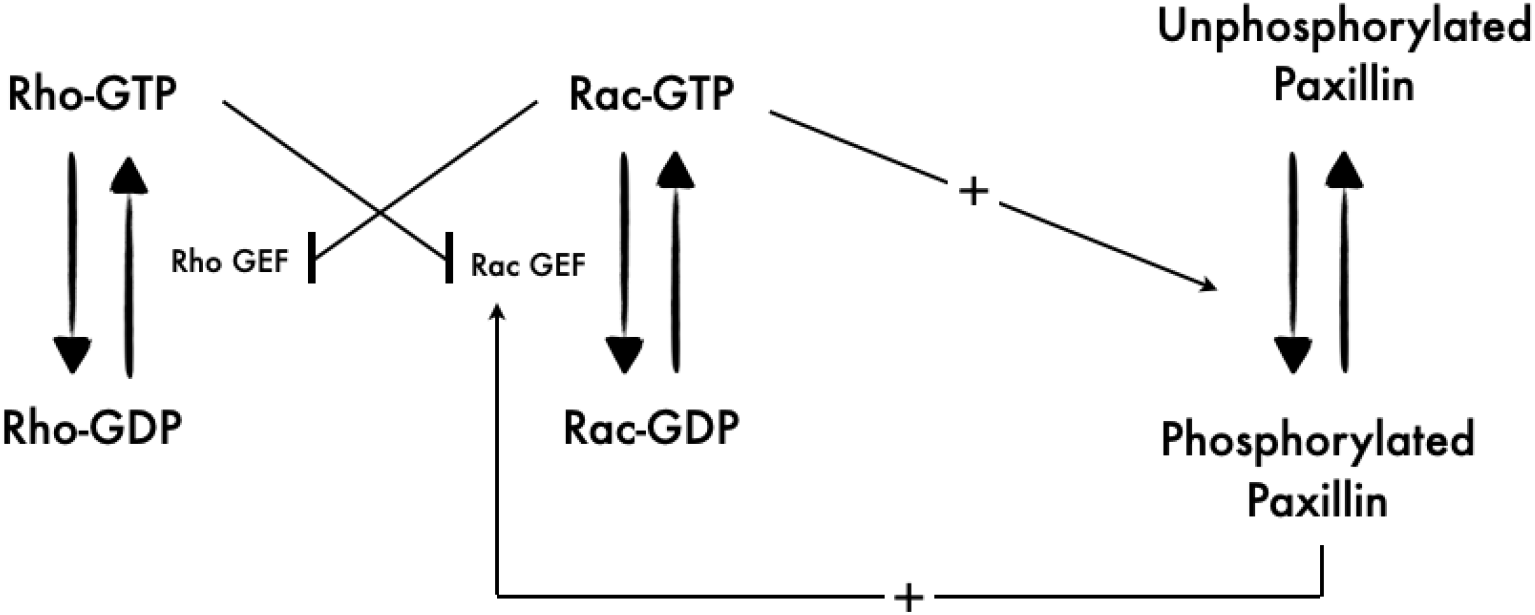
Schematic of the 6D model involving the Rac, Rho and paxillin regulatory network. Diagram of the signalling pathways of the 6D model showing the two proteins Rac and Rho cycling between inactive (GDP bound) and active (GTP bound) forms, and the active form of each protein inhibiting the activation of the other through GTPase-specific GEFs inactivation. It also shows how active Rac participates in paxillin phosphorylation at serine 273 (S273) residue which in turn promotes Rac activation in a positive feedback loop.

where *n* is the Hill coefficient, *L_ρ_* (*L_R_*) is the Rho-dependent (Rac-dependent) half maximum inhibition of Rac (Rho), *L_K_* represents the half-maximum phosphorylation of paxillin, *I_R_* is the basal Rac activation rate, *I_ρ_* is the Rho activation rate, *B* is the maximum paxillin phosphorylation rate, *δ_ρ_* (*δ_R_*) is the GAP-dependent Rho (Rac) inactivation rate, *δ_P_* is the paxillin dephosphorylation rate, *γ* is the ratio of total PAK to total Rac, *D_x_* (*x* = *R, R_i_, ρ, ρ_i_, P, P_i_*) is the diffusion coefficient of each molecular species, *K* is the scaled concentration of active PAK ([*PAK^*^*]), given by

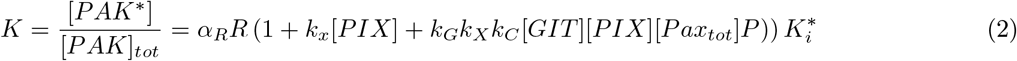

and 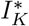 is the *GIT* − *PIX* − *PAK* complex-dependent Rac activation rate, given by

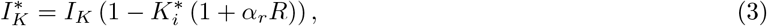

with

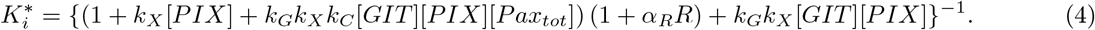

The parameters in Eqs. (2) and (3) represent the association constant for *PIX – PAK* binding (*k_X_*), the association constant for *GIT – PIX* binding (*k_G_*), the association constant for *Pax_p_ – GIT* binding (*k_C_*), the concentration of GIT and PIX ([*GIT*] and [*PIX*]) and the affinity constant for *PAK – RacGTP* binding (*α_R_*).

The first (second) terms in Eqs. (1a), (1c) and (1e) represent Rac, Rho activation (inactivation) and paxillin phosphorylation (unphosphorylation), respectively, whereas the third term represents the diffusion of these proteins. The terms in Eqs. (1b), (1d) and (1f) are defined similarly for the inactive/unphosphorylated form of these three proteins. Notice how the activation term of Rac (Rho) is a decreasing Hill function of Rho (Rac) to reflect the mutual inhibition exerted by the two proteins on each other, whereas the phosphorylation term of paxillin is an increasing function of Rac through *K*. The derivations of this highly nonlinear model are detailed in [26, 36]. For the complete list of parameter definitions and values, see Table 1.

**Table 1:** Parameter values associated with the original spatiotemporal model given by Eqs. (1).

| Parameter | Description | Value | Unit | References |
| --- | --- | --- | --- | --- |
| $I_R$ | Basal Rac activation rate | 0.0035 | $s^{-1}$ | [26] |
| $\delta_R$ | Rac inactivation rate | 0.025 | $s^{-1}$ | [26] |
| $L_\rho$ | Rho-dependent half-maximum inhibition of Rac | 0.34 | unitless | [26] |
| $I_\rho$ | Basal Rho activation rate | 0.016 | $s^{-1}$ | [26] |
| $\delta_\rho$ | Rho inactivation rate | 0.016 | $s^{-1}$ | [26] |
| $L_R$ | Rac-dependent half-maximum inhibition of Rho | 0.34 | unitless | [26] |
| $\gamma$ | Ratio of total PAK to total Rac | 0.3 | unitless | [26] |
| $\delta_P$ | Paxillin dephosphorylation rate | 0.00041 | $s^{-1}$ | [26] |
| $n$ | Hill coefficient | 4 | unitless | [26] |
| $L_K$ | Active PAK-dependent half-maximum activation of paxillin | 5.77 | unitless | [26] |
| $I_K$ | Additional Rac activation due to paxillin | 0.009 | $s^{-1}$ | [26] |
| $k_G$ | Association constant for GIT-PIX binding | 5.71 | $s^{-1}$ | [26] |
| $[GIT]$ | Concentration of GIT | 0.11 | $\mu M$ | [26] |
| $k_X$ | Association constant for PIX-PAK binding | 41.7 | $s^{-1}$ | [26] |
| $[PIX]$ | Concentration of PIX | 0.069 | $\mu M$ | [26] |
| $k_C$ | Association constant for $Pax_p$-GIT binding | 5 | $s^{-1}$ | [26] |
| $[Pax_{tot}]$ | Total concentration of paxillin | 2.3 | $\mu M$ | [26] |
| $\alpha_R$ | Affinity constant of PAK-RacGTP binding | 15 | unitless | [26] |

## 3 Model reduction and simplification into 1D ODE model

Due to the high dimensionality of the original 6D spatiotemporal PDE model (1) and the difficulty in interpreting its underlying dynamics, Tang *et al* [26] simplified the model and reduced its dimensionality. This was done by first ignoring the diffusion terms, turning the PDE model into an ordinary differential equation (ODE) model, followed by assuming that protein biosynthesis in the cell is slow enough, implying that the total amount of each protein is conserved (i.e., constant). This produced the following system of ODEs

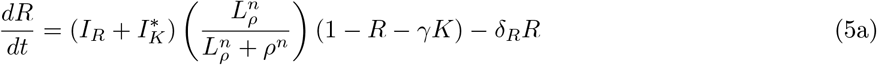

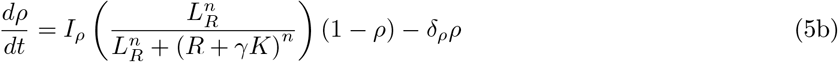

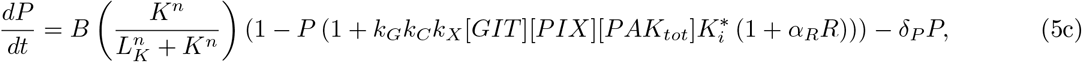

where *R_i_* = 1 − *R*, *ρ_i_* = 1 − *ρ* and 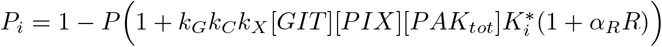.

To further simplify this ODE model without modifying its underlying dynamics, several simplifying steps have been implemented here to reduce its dimensionality. In the first step, we apply linear regression on 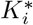, defined by Eq. (4), using the values of paxillin at steady state; this generates the following simplified expression

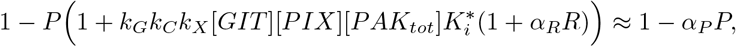

where *α_P_* is a parameter (see Table 2). Second, since the dynamics of active Rho (*ρ*) and phosphorylated paxillin (*P*) occur at a faster timescale compared to active Rac (*R*), one can use quasi-steady state approximation (QSSA) to express *ρ* and *P* as functions of *R*. This is done by setting Eqs. (5b) and (5c) to zero to obtain

**Table 2:** Parameter values associated with the 2D and 3D ODE models defined by by Eqs. (9) and (10), respectively.

| Parameter | Description | Value | Unit |
| --- | --- | --- | --- |
| $\alpha_P$ | Linearization coefficient in paxillin activation | 2.7 | unitless |
| $\epsilon$ | Time constant for $B$ | 0.01 | $s^{-1}$ |
| $B_r$ | Resting state of $B$ | 10 | unitless |
| $k_B$ | Recovery rate of $B$ back to its resting state | 0.04 | unitless |
| $\gamma_R$ | Strength of $R$ feedback onto $B$ | 8.6956 | $s^{-1}$ |
| $\eta, \epsilon_B$ | Parameters that guarantee the positivity of $B$ | $10^4, 10^{-4}$ | unitless |
| $\epsilon_L$ | Time constant of $k_B$ | $10^{-5}$ | $s^{-1}$ |
| $\gamma_K$ | Source term | 0.15 | unitless |

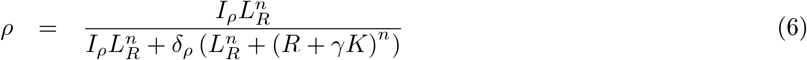

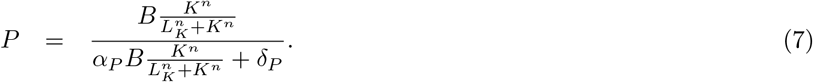

Finally, at steady state, *K* defined by Eq. (2) can be approximated (through fitting) using an expression that purely depends on *R*. This fitted expression is given by

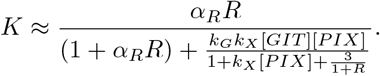

The same can be done for 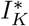, defined by Eq. (3), to obtain a simplified expression in terms of *R* only, given by

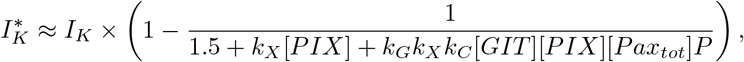

where *P* is at steady state defined by Eq. (7).

Taking all these approximations into consideration, including QSSA, produces a 1D ODE model for *R*, given by

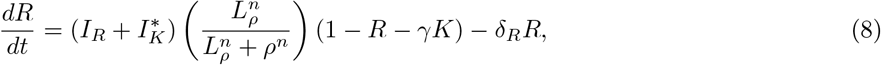

where *ρ*, *K* and 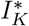 are defined by Eqs. (6), (2) and (3), respectively. This new 1D ODE model is in a closed form as it purely depends on *R*.

The bistability in *R* with respect to maximum phosphorylation rate of paxillin *B* was an important feature of the original ODE model given by Eqs. (5) [26]. This is further confirmed here by plotting the bifurcation diagram of *R* with respect to *B* (Fig. 2A), producing a bistable switch comprised of two branches of stable equilibria (solid lines) linked together by a branch of unstable equilibria (dashed line) at two saddle-node bifurcations. To determine if such bistable switch is preserved by the simplified 1D ODE model, we plot the same type of bifurcation diagram for *R* in terms of *B* but use Eq. (8). Doing so produces a bistable switch whose shape and range of hysteresis are similar to that obtained by the original ODE model (Fig. 2B). In both models, bistablity allows for the switching between the uninduced (low active Rac) and induced (high active Rac) states to occur at specific thresholds in *B* (determined by the saddle-node bifurcations), outside of which the system becomes monostable. These results thus demonstrate that the simplified 1D ODE model, defined by Eq. (8), preserves the key dynamic features of the original ODE model, defined by Eq. (5), and can thus be used to analyse motility.

**Figure 2:**
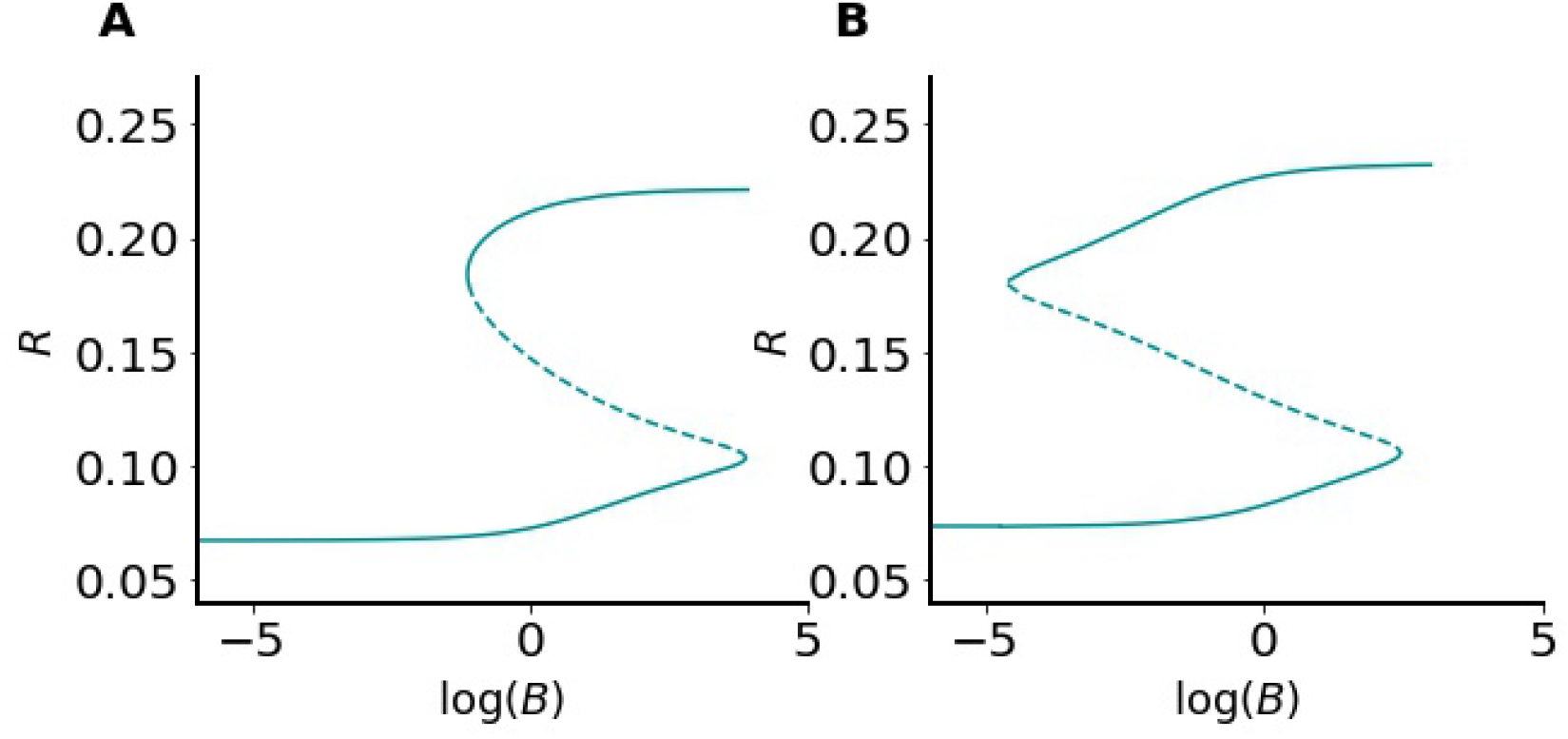
Bistable switch generated by the Rac, Rho and paxillin signaling network. The bifurcation diagram of *R* with respect to *B* according to the (A) 6D, and (B) 1D models defined by Eqs. (1) and (8), respectively, when diffusion is ignored. Both diagrams exhibit bistability between the uninduced (low active Rac) and induced (elevated active Rac) states in an almost identical fashion. Solid lines: branches of stable equilibria; dashed lines: branches of unstable equilibria (or saddle fixed p oints). Solid lines merge with dashed lines at saddle-node bifurcations.

## 4 Two-dimensional (2D) ODE model with one fast and one slow variable

Studying the impact of bistability on cellular polarization has been previously investigated in detail in past modeling studies [37, 38]. Because CHO-K1 cells are known to exhibit much more complex migration patterns (manifested as quick increases and decreases in protrusions, indicating variations in the Rac activity) [31], we have decided to increase the complexity of the 1D ODE model, given by Eq. (8), to allow the modified model to display a whole set of new and interesting dynamics, including oscillations. The goal is to explore how such new complex dynamics can impact motility.

With the evidence suggesting that there are potential feedbacks from Rac to maximum phosphorylation rate *B*, the 1D ODE model is modified here accordingly. This is done by augmenting the 1D ODE model with one additional auxiliary and phenomenological equation that reflects the slow dynamics of paxillin phosphorylation. More specifically, we have turned the parameter *B* into a slow variable, given by

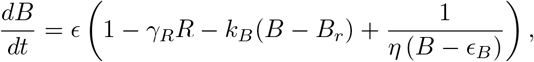

where 0 < *ϵ* ≪ 1 is the time constant, *B_r_* > 0 is the resting state of *B*, *k_B_* > 0 is the recovery rate to the resting state and *ϵ_B_, η* > 0 are parameters. The resulting 2D ODE model becomes

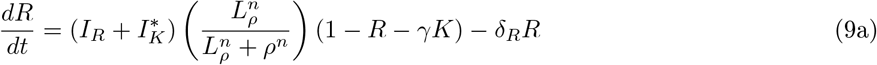

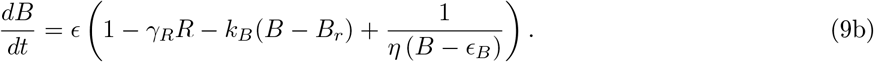

This new 2D ODE model is designed to have *B* vary very slowly compared to *R*. This means that *R* can always converge to a steady state, if it exists, before any substantial variation in *B*. The slow dynamics of *B* is depicted here by *ϵ*. Two important features of this new model include the absence of feedback from *B* to active Rac *R* (i.e., the model is feedforward with *B* receiving input from *R* but not vice versa), and the impossibility for *B* to be negative due to the term 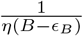 which acts as a barrier.

Equations 9, possess one slow and one fast variable. As before, this model can exhibit bistability for certain parameter values. Indeed, depending on the parameters that govern the variation in *B* (e.g., *k_B_* and *γ_R_*), the system can either have one or two co-existing stable steady states, when the *B*-nullcline crosses the “generalized” *R*-nullcline at stationary points where *R* is stable (i.e., the lower and upper stable branches of the generalized *R*-nullcline, see Fig. 3A, B and C). The real interesting dynamic feature of this new model, however, is that it can also exhibit ROs; this occurs when it is in a configuration in which the *B*-nullcline intersects the generalized *R*-nullcline in the middle unstable branch (Fig. 4A). This would allow the 2D ODE model to exhibit relaxation oscillations (ROs) along the hysteresis cycle that involves the slow evolution of trajectories along the stable branches of the generalized *R*-nullcline, because *B* is slow, and the fast jumps between the two stable branches at the saddle-nodes, because *R* is fast (Fig. 4B). Having the *B*-nullcline separating the phase-space into two regions: one above the *B*-nullcline with *dB/dt* < 0 (*B* is decreasing) and another below the *B*-nullcline with *dB/dt* > 0 (*B* is increasing), causes the trajectories to move to the left (right) below (above) the *B*-nullcline until they reach the saddle-nodes. Thus for such oscillations to occur, *ϵ* must be very small and the two nullclines must intersect only once on the middle unstable branch of the cubic-shaped generalized *R*-nullcline (Fig. 4).

**Figure 3:**
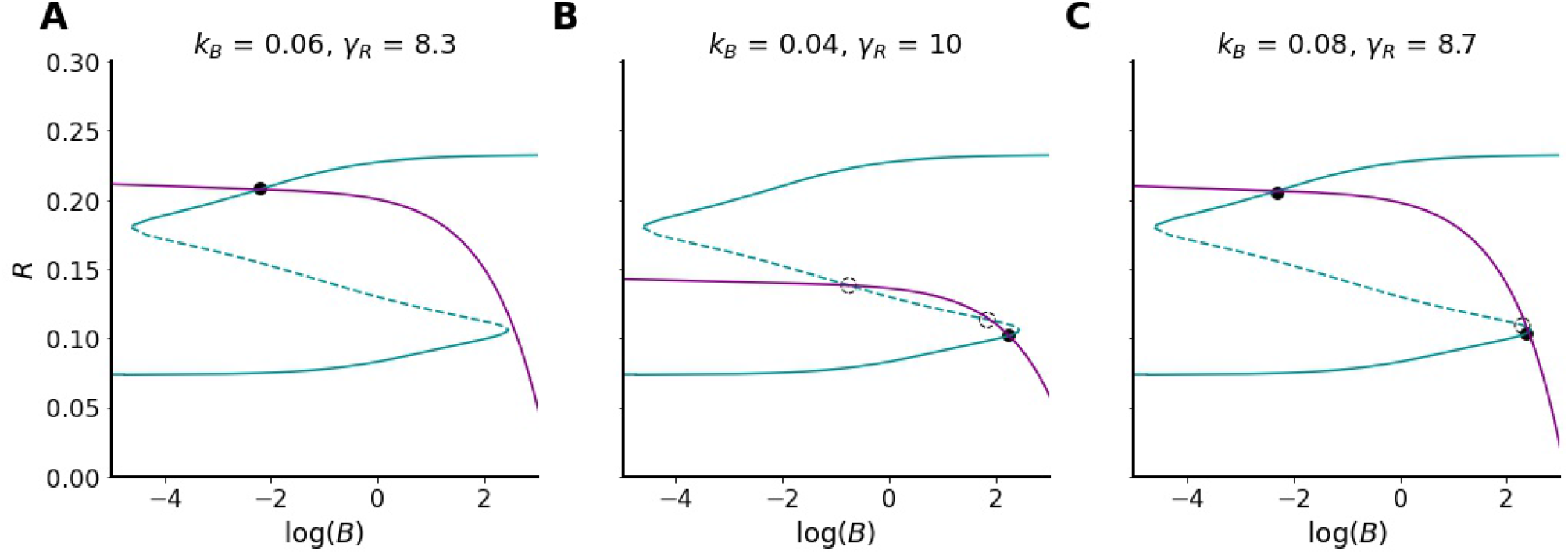
Three possible configurations for the steady states of the 2D ODE model, given by Eqs. 9, lying at the intersection of the *B*-nullcline (purple) and generalized *R*-nullcline (blue) obtained when *k_B_* and *γ_R_* are varied but no change is made to the remaining parameter values listed in Tables 1 and 2. In the first two cases, one stable steady state, lying either on the (A) upper or (B) lower stable branch of the generalized *R*-nullcline is formed between the two nullclines. In the last case, (C) two stable steady states, lying on the upper and lower stable branches of the generalized *R*-nullcline, are formed. Black solid (empty) circles represent stable (unstable) steady states.

**Figure 4:**
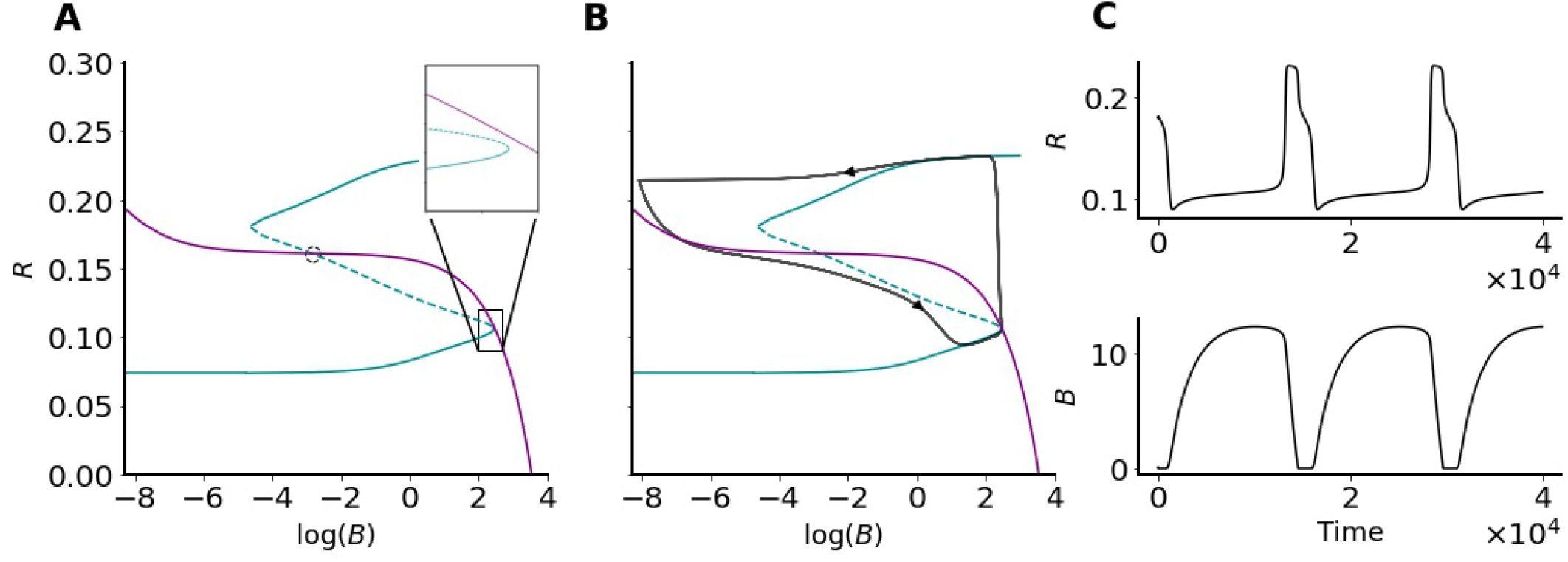
A new configuration for the steady state of the 2D ODE model, given by Eqs. 9, lying at the intersection of the *B*-nullcline (purple) and generalized *R*-nullcline (blue) obtained at default parameter values listed in Tables 1 and 2. In this case, (A) only one intersection occurs on the middle unstable branch of the cubic generalized *R*-nullcline, forming an unstable steady state (open circle). The inset in (A) is a magnification of the box surrounding the bottom saddle-node bifurcation. (B) Stable periodic orbit in the form of ROs (black) is formed around the unstable steady state. The cause for the apparent stretching of the ROs far from the generalized *R*-nullcline in (A, B) is the use of the logarithmic scale, which considerably expands any small deviation near zero. (C) Time series simulations of *R* (top) and *B* (bottom) showing oscillations.

With these ROs, one would expect (as shown later) to obtain a Hopf bifurcation when varying the parameter(s) related to *B*, or more specifically those that dictate the position of the *B*-nullcline (e.g., *k_B_*). Indeed, if this parameter variation causes the configuration of the nullclines in Fig. 3A, where the two nullclines intersect at a stable steady state in the upper branch of the generalized *R*-nullcline, to transition to the one in Fig. 4A, where the two nullclines intersect at an unstable steady state in the middle branch of the generalized *R*-nullcline, a shift in dynamics from quiescent to oscillatory behaviour would occur right when the intersection of the two nullclines crosses the upper saddle-node bifurcation during this transition. This oscillatory behaviour is eventually manifested as a RO (Fig. 4B).

Interestingly, during the transition from quiescent to the ROs obtained while varying the slope of the *B*-nullcline (through the parameter *k_B_*), another very interesting dynamics is also observed within a very small parameter regime (Fig. 5A). In this third regime, trajectories exhibit very small amplitude oscillations (SAOs) in both variables: *R* and *B* (Fig. 5B). Such oscillations are formed as soon as the stable steady state disappears during the decrease in the slope of the *B*-nullcline and a new unstable steady state is formed in the middle branch of the generalized *R*-nullcline. Trajectories, in this case, initially get attracted to the lower stable branch of the generalized *R*-nullcline causing *R* to jump from the upper branch at the saddle-node bifurcation; during this jump, however, the fast variable crosses the *B*-nullcline, causing the slow variable *B* to increase again. When *B* reaches a higher value, it brings the system back to the right of the upper saddle-node bifurcation. Meanwhile, the decrease in *R* is insufficient to bring the trajectories all the way down to the lower branch, causing them to interrupt their jumps and to return back to the upper branch of the generalized *R*-nullcline (because of the increase in *B*). When this happens, the same phenomenon repeats itself again, causing both *R* and *B* to generate SAOs (Fig. 5B).

**Figure 5:**
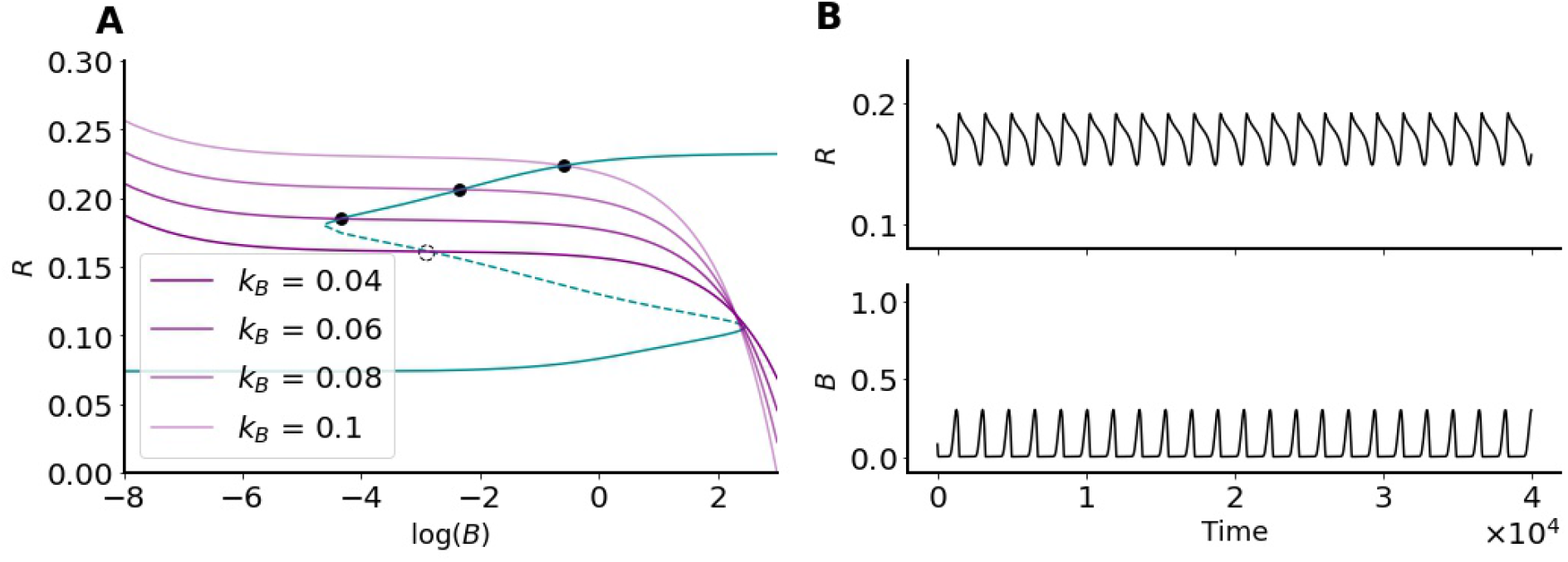
The 2D ODE model, given by Eqs. 9, can exhibit SAOs when varying *k_B_* due to canards. (A) Gradually decreasing the slope of the *B*-nullcline through *k_B_*, the steady state at the intersection of the generalized *R*-nullcline (blue) and the *B*-nullcline (shades of purple) transitions from being stable (lying on the upper branch of the generalized *R*-nullcline) to being unstable (lying on the middle branch of the generalized *R*-nullcline). (B) Time series simulations of SAOs displayed by *R* (top) and *B* (bottom) formed right after the *B*-nullcline shifts its intersection with the generalized *R*-nullcline to the middle branch.

### 4.1 Three-dimensional (3D) ODE model with one fast, one slow and one very slow variable

The SAOs highlighted in the previous section (Fig. 5) occur between the upper stable branch of the generalized *R*-nullcline and its unstable branch, in the neighbourhood of the upper saddle-node. Further investigation of the 2D ODE model of Eqs. (9) shows that for the trajectories to go back and forth between the quiescent configuration of Fig. 3A to the RO configuration of Fig. 4, one needs to periodically decrease and increase the slope the *B*-nullcline sufficiently enough. This can be done by targeting the the recovery rate parameter *k_B_* which defines the slope of the *B*-nullcline. By dynamically varying *k_B_*, the system can travel through the whole spectrum of these regimes highlighted earlier, producing SAOs in combination with ROs in the form of large amplitude oscillations (LAOs). This combination of different amplitude oscillations is called mixed modes oscillations (MMOs).

Keeping that in mind, we have expanded the 2D ODE model into a 3D semi-phenomenological ODE model based on the scheme of Fig. 6 to produce MMOs. The model possesses three different timescales: fast, slow and very slow, to allow for the aforementioned patterns of activity to arise, including MMOs that combine slow large amplitude with fast small amplitude oscillations within one cycle. The fast variables in this new model is *R* whereas the slow and the very slow variables are the two parameters – turned into variables – *B* and *k_B_*, respectively. The 3D ODE model is given by

**Figure 6:**
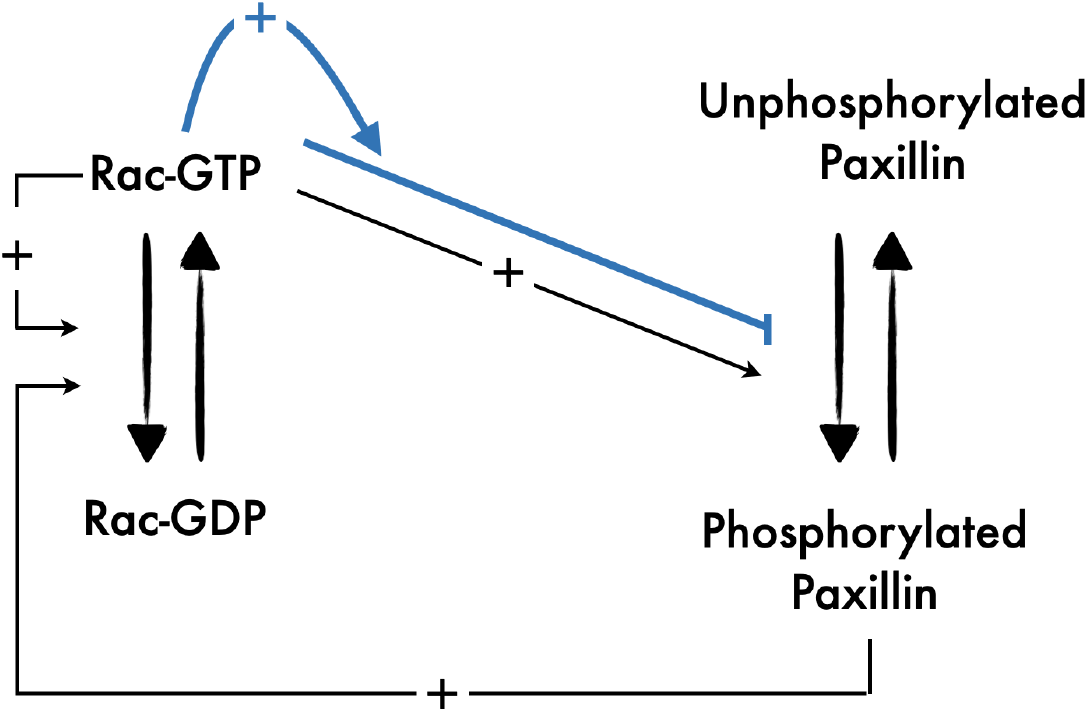
Schematic of the 3D ODE model involving Rac and paxillin in a simplified protein network. Diagram of the signalling pathways included in the 3D ODE model showing Rac cycling between inactive and active forms and the indirect positive auto-feedback (via Rho) on itself through RacGEF. It also shows the positive and negative (blue pathway) feedbacks on paxillin phosphorylation at a fast and a slow timescale, respectively, as well as the indirect upregulation of Rac activation through RacGEF by phosphorylated paxillin. Blue pathway: refinement of the original model in [26].

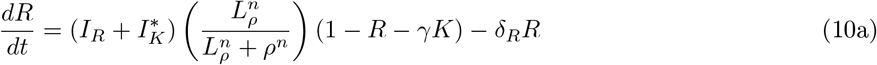

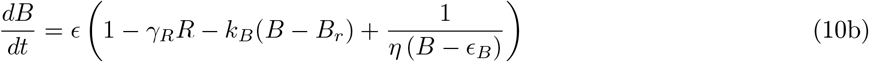

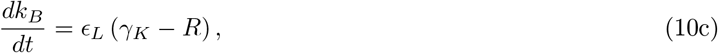

where 0 < *ϵ_L_* ≪ *ϵ* < 1 is the time constant and *γ_K_* > 0 is a parameter chosen to be between the steady states of *R* defined by the generalized *R*-nullcline. The equations for *R* and *B* are identical to these in Eqs. (9).

The 3D ODE model of Eqs. (10) is once again feedforward. The new auxiliary equation for recovery variable *k_B_* possesses the slowest timescale defined by *ϵ_L_*, receives input from *R* in a feedforward manner and allows the system to exhibit oscillations of varying amplitudes and periods. These oscillations are manifested as MMOs in *R* (Fig. 7).

**Figure 7:**
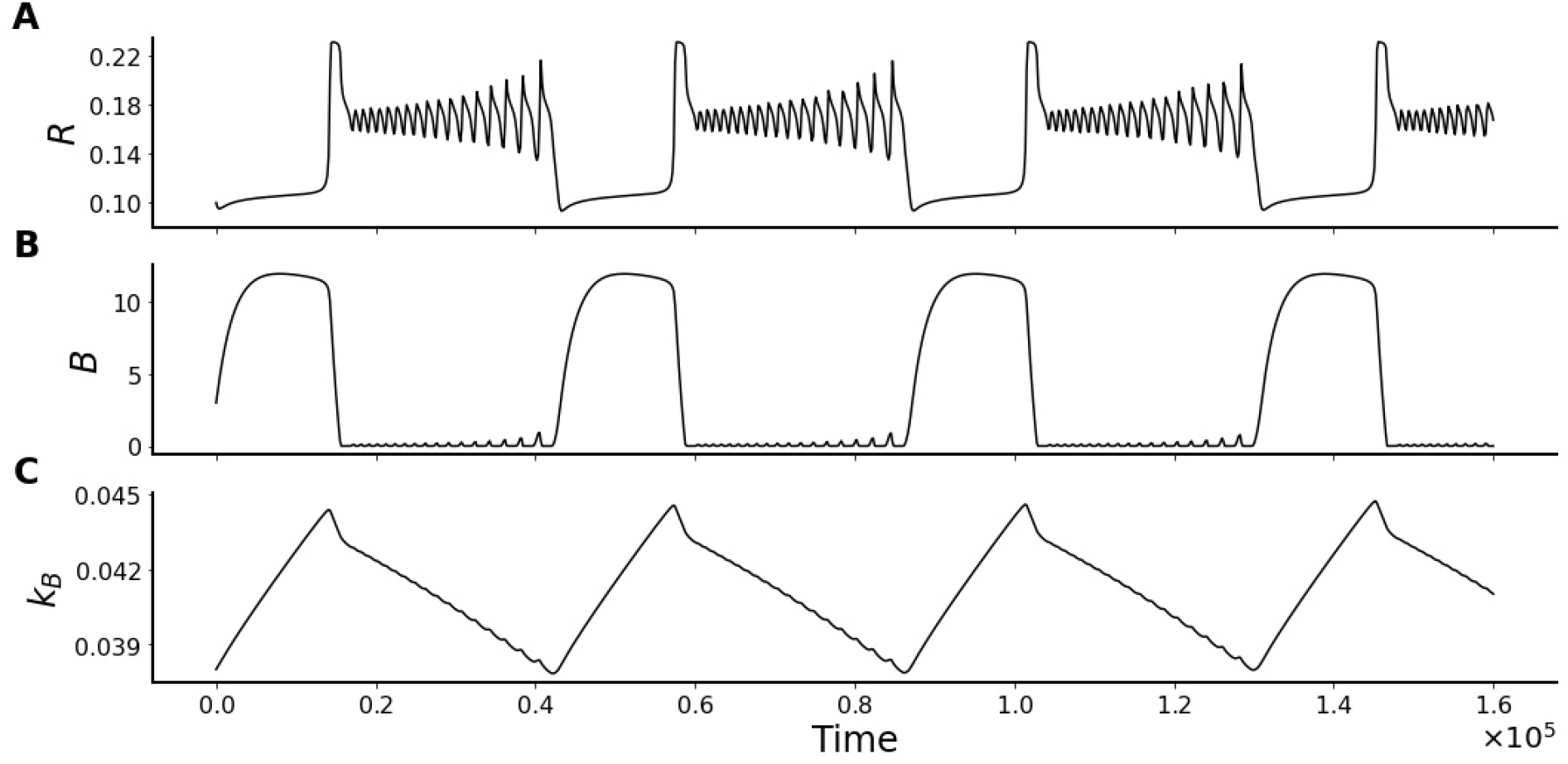
MMOs generated by the 3D ODE model defined by Eqs. (10). Time course simulations of (A) *R*, (B) *B* and (C) *k_B_* showing oscillations in each variable with *R* displaying very pronounced MMOs.

### 4.2 Slow-fast analysis of the 3D ODE model

The MMOs detected in the 3D ODE model of Eqs. (10) (Fig. 7) indicate that there is a quick and exponential increase in the amplitude of these oscillations through a canard explosion. This classical canard phenomenon explains the very fast transition from a small amplitude oscillations (SAO), via canard cycles, to a large amplitude oscillation (LAO), after a Hopf bifurcation. Such very fast transition happens within an exponentially small range of a control parameter, making it difficult to detect. Still, the peculiar shape of the generalized *R*-nullcline (i.e., the critical manifold of the 3D ODE model) makes our system very prone to developing these canard explosions; indeed, these types of oscillations are always encountered when designing the model and setting different formulations for the *B* and *k_B_* equations.

Desroches *et al* (2012) [32] provided a detailed summary of the local mechanisms that give rise to MMOs. As acknowledged in this study, different configurations that can produce MMOs have been widely studied in the presence of folded singularities in the critical manifold. According to this scenario, the simplest form for a system to exhibit MMOs is to have a three-dimensional model with one fast and two slow variables. By investigating the dynamics of the reduced problem, obtained by setting the fast variable to steady state, one could find folded singularities in the critical/slow manifold of the problem and thus explain the presence of MMOs. However, in some cases, there is no folding in the critical manifold, and therefore another less common approach must be applied, where the system is viewed as consisting of two fast variables and one slow one. Here, we adopt this approach with the 3D ODE mode, given by Eqs. (10), by taking *R* and *B* to be fast and *k_B_* to be slow. To simplify the analysis, we let

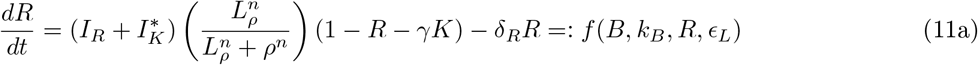

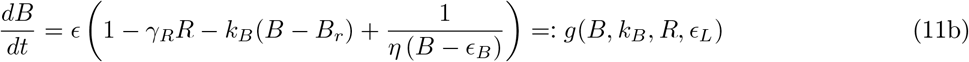

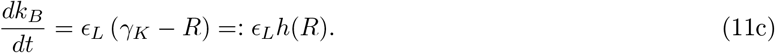

Based on this, we can define the layer problem to be

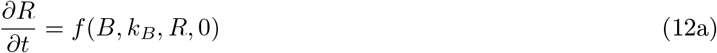

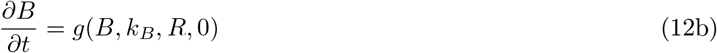

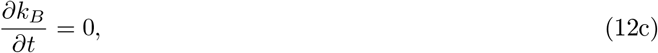

and the the reduced problem, obtained by transforming the fast time scale *t* into a slow time scale *T* = *ϵ_L_t*, to be

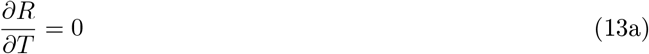

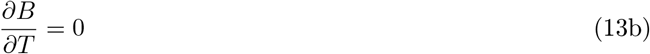

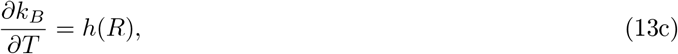

where *ϵ_L_* is set to zero in both problems defined by Eqs. (12) and (13). According to this formalism, the critical manifold is defined to be 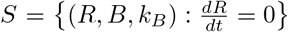, which is equivalent to the null-surface of the fast variable *R*.

With these two problems introduced, we can now plot the bifurcation diagrams of the fast variables with respect to *k_B_* as defined by the layer problem of Eqs. (12) (Fig. 8). Our results reveal that both variables of the fast sub-system, namely, *R* (Fig. 8A) and *B* (Fig. 8B), undergo a supercritical Hopf bifurcation when decreasing *k_B_* at *k_BHopf_* ≈ 0.048. Two envelopes of stable periodic orbits representing the maximum and minimum of the oscillations emanate from the Hopf bifurcation. The amplitude of the oscillations increases exponentially fast in the form of a canard explosion until these envelops eventually plateau starting from approximately *k_BLAO_* ≈ 0.038 (estimated using AUTO-07p [39]) towards smaller values of *k_B_*. The periodic orbits associated with canard explosion seen in the bifurcation diagrams of *R* and *B* (Fig. 8A and B, respectively) are ROs that partially contribute to the formation of the LAOs in the MMOs. Due to the fact that these oscillations appear at *k_BLAO_ < k_BHopf_*, the Hopf bifurcation in this case is referred to as delayed Hopf. In our subsequent analysis, we use the phrase “active phase” of the system to refer to the configuration when *k_B_* is between *k_BLAO_* and *k_BHopf_*.

**Figure 8:**
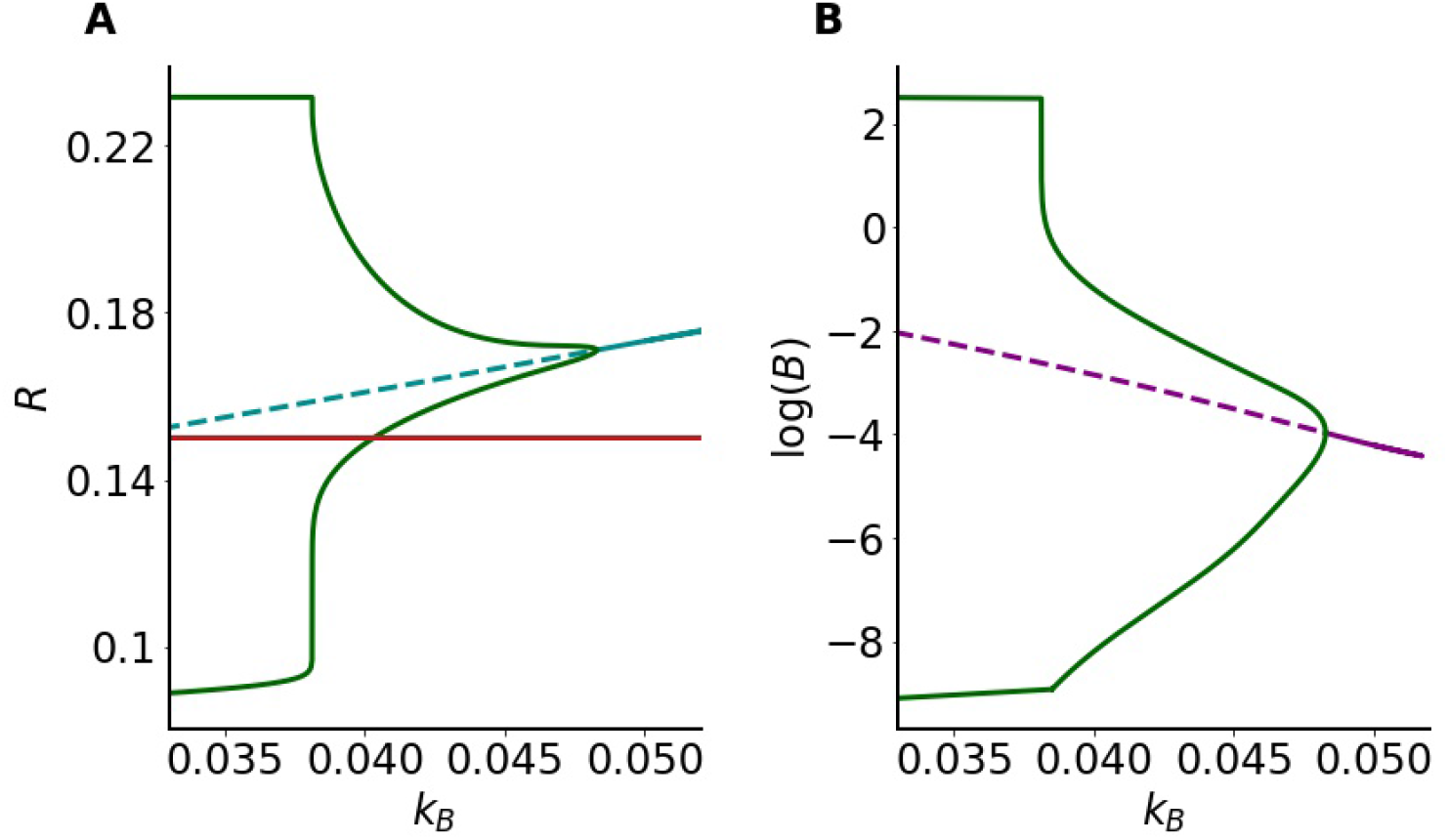
Slow-fast analysis of the 3D ODE model defined by Eqs. (10). Bifurcation diagram of (A) *R* and (B) *B* with respect to *k_B_* as defined by the layer problem 12. Solid (dashed) blue and purple lines represent stable (unstable) equilibria. In both panels, the change in stability (i.e., the switch from solid to dashed line) occurs at a supercritical Hopf bifurcation at which two envelopes of stable periodic orbits (solid green), representing the maximum and minimum of the oscillations, emerge. Red line in Panel A is the *k_B_*-nullcline superimposed on the bifurcation diagram.

To further investigate how SAOs and LAOs form MMOs, we now turn our attention to the reduced problem, given by System 13. By superimposing the nullcline of *k_B_* on the bifurcation diagram of *R* with respect to *k_B_* (Fig. 8A), we can see that the dynamic of the full 3D ODE model is dictated by how long trajectories spend time below or above the *k_B_*-nullcline. Indeed, if the average value of *R*, calculated over an entire SAO cycle, is bigger than *γ_K_*, then *k_B_* will decrease (since the average *dk_B_/dt* < 0), causing the trajectory to move to the left in Fig. 8. If the opposite happens, then the trajectory moves to the right. Such mechanism is similar to that seen in parabolic bursting [40].

To verify if this is the mechanism underlying MMOs, one cycle of the solution trajectory of the full 3D ODE model, given by Eqs. (10), is superimposed onto the bifurcation diagram of *R* with respect to *k_B_* (Fig. 9A). Doing so shows that the trajectory initially moves to the left while exhibiting SAOs delimited by the two envelopes of the periodic orbits to the right of the canard explosion because it spends more time above the *k_B_*-nullcline. Thus, the number of spikes during the active phase is directly linked to the time constant of *k_B_*, because the time the system spends in the active phase depends on how fast *k_B_* decreases. However, when the trajectory crosses *k_BLAO_* and enters the relaxation oscillation regime, it starts spending more time below the *k_B_*-nullcline causing it to travel to the right (recall that we have *R_low_ < γ_K_ < R_high_*).

**Figure 9:**
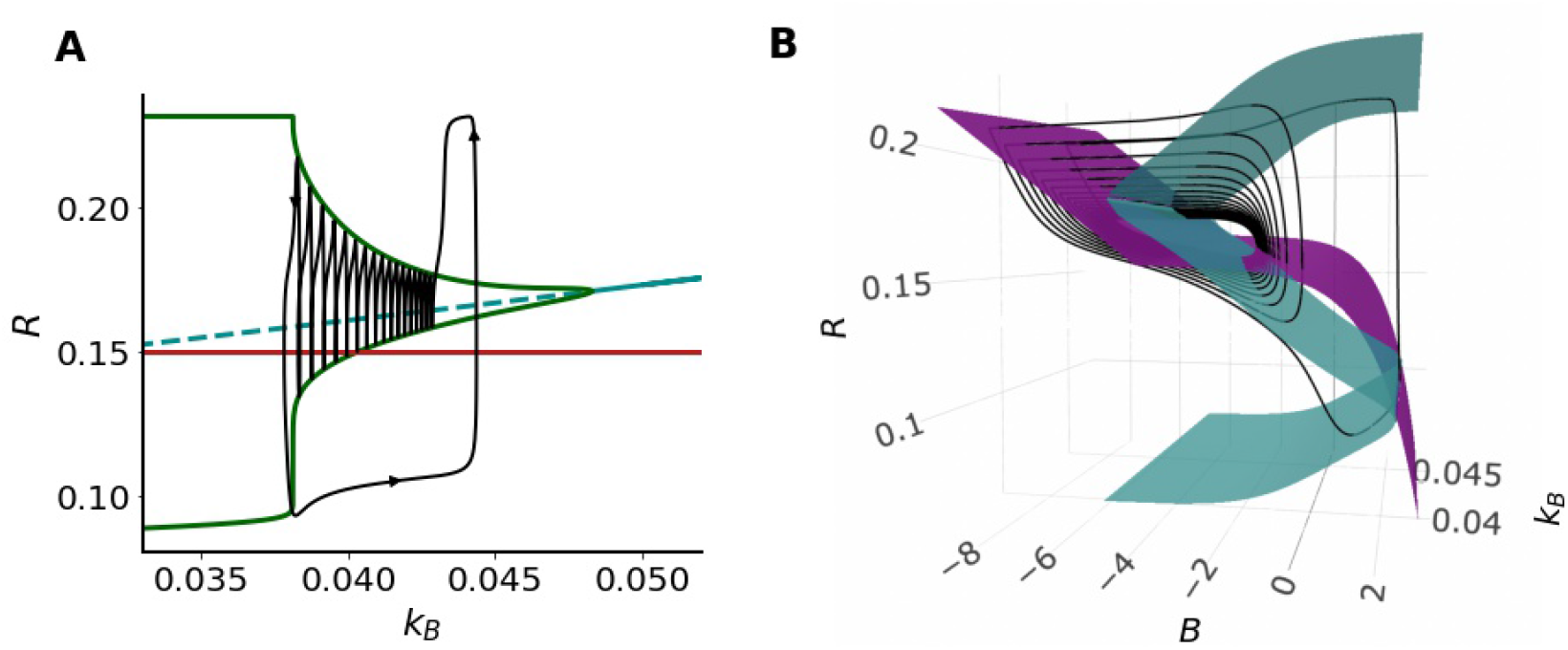
Underlying dynamics of MMOs produced by the 3D ODE model defined by Eqs. (10). (A) Solution trajectory (black line) superimposed in the bifurcation diagram of *R* with respect to *k_B_* previously plotted in Fig 8. (B) The critical manifold *S* (blue) along with the *B*-nullsurface (purple) showing how they govern the dynamics of a solution trajectory (black) over time.

Notice here that, according to the configuration of Fig. 9A, the trajectory does not spend too much time in the RO regime, as it leaves very soon after by moving to the right away from the periodic envelopes. The contribution of ROs (through the canard explosion) to the MMOs is therefore limited. Interestingly, when the trajectory moves to the right, it peculiarly does not follow a specific manifold and suddenly changes direction upward by forming a very prominent LAO. The configuration of Fig. 9A is thus not sufficient to explain all the dynamics.

To explain the underlying dynamics of this very last component of the solution trajectory showing the prominent LAO, it is necessary to expand our slow-fast analysis to include *B*. This is done in Fig. 9B, where we plot the critical manifold of the layer problem, given by Eqs. (12), and the *B*-nullsurface along with one cycle from the solution trajectory of the full 3D ODE model to view how it evolves with respect to both of these surfaces. Our results reveal that towards the end of the last SAO (which is becoming a LAO), the trajectory reaches the lower stable sheet of the critical manifold. When this happens, the trajectory ends up being below the *B*-nullsurface in a region where *dB/dt* > 0, causing it to travel right along this stable sheet until it reaches the fold (the saddle-node); this produces the lower plateau part of the trajectory in Fig. 9A right after leaving the oscillatory regime. At the fold, the trajectory then jumps up to the upper stable sheet of the critical manifold of Fig. 9B (because *R* is fast), crossing the *B*-nullsurface into a region where *dB/dt* < 0, causing the trajectory to travel to the left briefly along the upper sheet until it crosses the *B*-nullsurface all over again (Fig. 9B). Because *B* has slow time scale, trajectory then moves downward and slightly to the left along the *B*-nullsurface until it enters the oscillatory regime determined by *k_B_*. The full cycle of jumping up, and down with a short plateau in between form the LAO seen in Fig. 9A.

### 4.3 Spatiotemporal dynamic in the presence of diffusion

By reintroducing diffusion into the *R* equation of the 1D and 3D ODE models, given by Eqs.(8) and (10), respectively, one can investigate the spatiotemporal dynamic of Rac according to these models. In order to do this in the context of cell motility, however, one also needs to reintroduce the equation that govern the dynamics of inactive Rac (*R_i_*). This produces, as a result, two spatiotemporal models that we label as the 2D and 4D PDE models, respectively.

#### 4.3.1 Wave-pinning in the 2D PDE model

A hallmark of cell motility and cell polarization is the formation of a stable inhomogeneous pattern in molecular concentrations between the front and the rear of a polarized cell [41]. Such a pattern has been previously referred to as wave-pinning [38, 42]; it is a spatiotemporal phenomenon that describes the propagation of a front of higher protein concentration, originating from a perturbation of a homogeneous low steady state, and the stabilization of the front, creating a restricted region with high concentration of an active protein at steady state. It is an inherent property of some reaction-diffusion systems possessing bistability, and relies on two important biophysical assumptions, namely, that matter is conserved and that there is a difference in diffusion coefficients between the chemical species under consideration [38, 42].

As an example, let us consider the 2D PDE model, given by

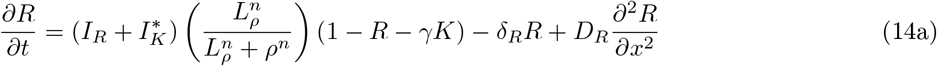

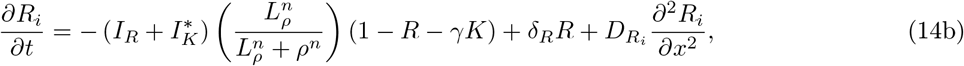

where *ρ* is defined by Eq. (6). This model depends purely on active (*R*) and inactive (*R_i_*) Rac in a closed form, is conserved and possesses differences in diffusion coefficients between its two molecular species: *R* and *R_i_*. In other words, it has all the ingredients necessary to produce wave-pinning.

To verify this, the model is simulated over time on a one-dimensional spatial domain by initiating it from non-homogeneous initial condition (Fig. 10). A step function with *R* = 0.083 for *x* ≤ 30 and *R* = 0.2269 for *x* > 30 (which correspond, respectively, to the stable low and high equilibria of R in the 1D ODE model) is used as an initial condition. By doing so, we obtain the wave-pinning phenomenon comprised of elevated *R* in the "front” and low *R* in the “back” (Fig. 10A). This spatiotemporal phenomenon remained stable at steady state due to the low diffusion coefficient of *R*. In contrast, the high diffusion coefficient of *R_i_* prevented this from happening, leading *R_i_* to become homogeneous at steady state (Fig. 10B). This result thus confirms that the reduced 2D PDE model of Eqs. 14 can produce wave-pinning in Rac in a manner similar to that seen with the 6D PDE model [26].

**Figure 10:**
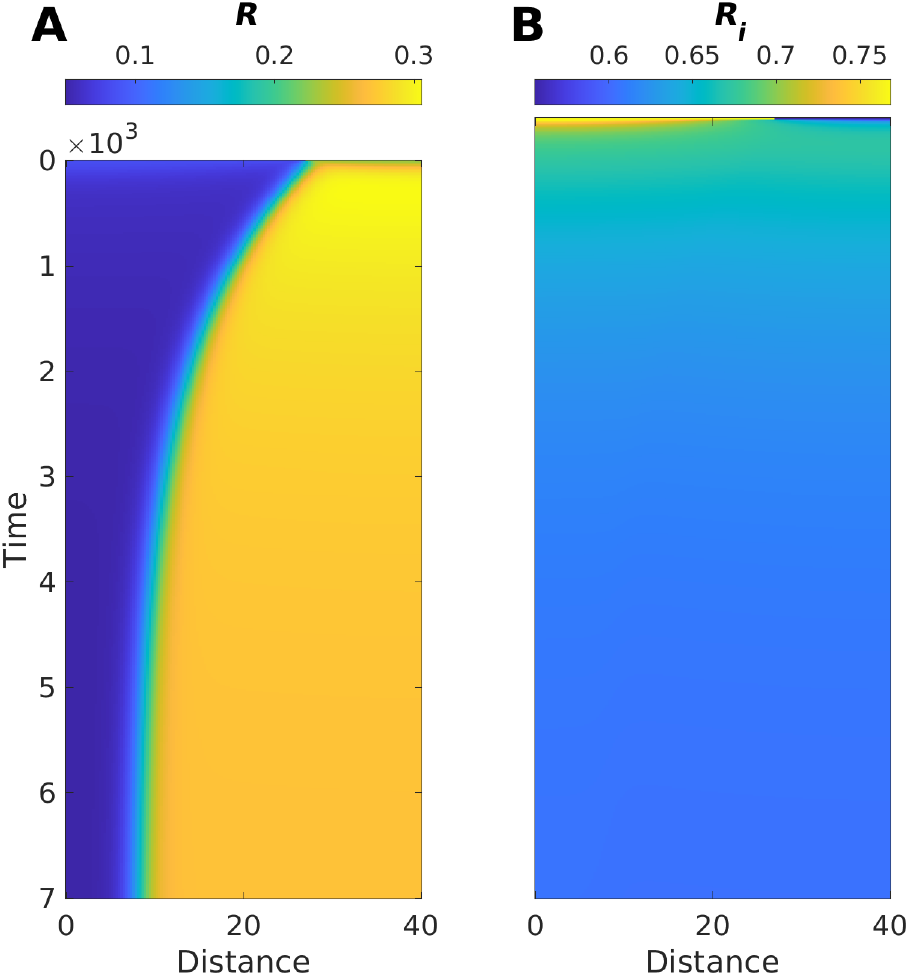
Wave-pinning in a one-dimensional spatial domain generated by the 2D PDE model, given by Eqs. (14). Kymographs of (A) *R* and (B) *R_i_* when a step-function is used as an initial condition, with zero-flux boundary conditions. The kymographs are color-coded according to the color-map in each panel. *R_i_* is homogeneous, having 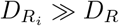, whereas *R* exhibits a front that eventually gets pinned. Longer simulations indicate that the front stays indefinitely in a fixed position, preventing the homogenization of the domain in terms of *R*. Time and distance are dimensionless.

#### 4.3.2 Spatiotemporal dynamics of the 4D PDE model

To explore how MMOs affect the spatiotemporal patterns generated by Rac, we turn our attention now to the 4D PDE given by

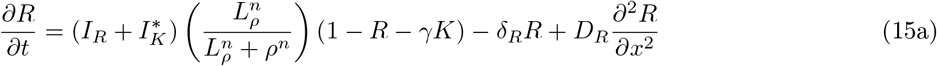

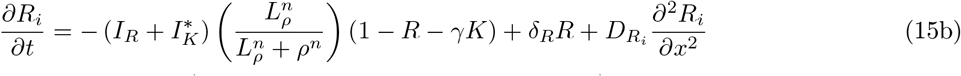

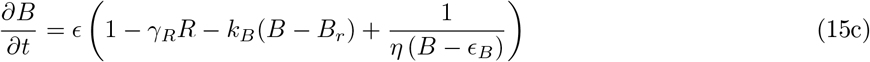

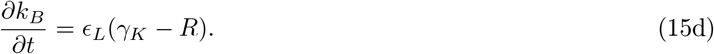

The numerical instability of this model makes it quite difficult to simulate it over time on a one-dimensional domain. To overcome this obstacle, we revert *k_B_* to being a parameter and analyze the dynamics of the resulting 3D PDE model in two different parameter regimes associated with *k_B_*: one that produces ROs and another that produces SAOs (as defined by the 2D ODE model of Eqs. (9)).

Our results show that, by simulating this 3D PDE model of active (*R*), and inactive (*R_i_*) Rac and *B* in the RO regime, we obtain intermittent polarization that switches orientation back and forth as time evolves (Fig. 11A). In the SAO regime, however, the entire domain shows small ripples that are present in all variables, especially in *R* (Fig. 11B). Interestingly, the switching in the level of the three variables in the SAO regime at the boundary of the domain is less pronounced than those seen in the RO regime (compare Fig. 11A to B), but the scaled concentrations across the domain are more heterogeneous.

**Figure 11:**
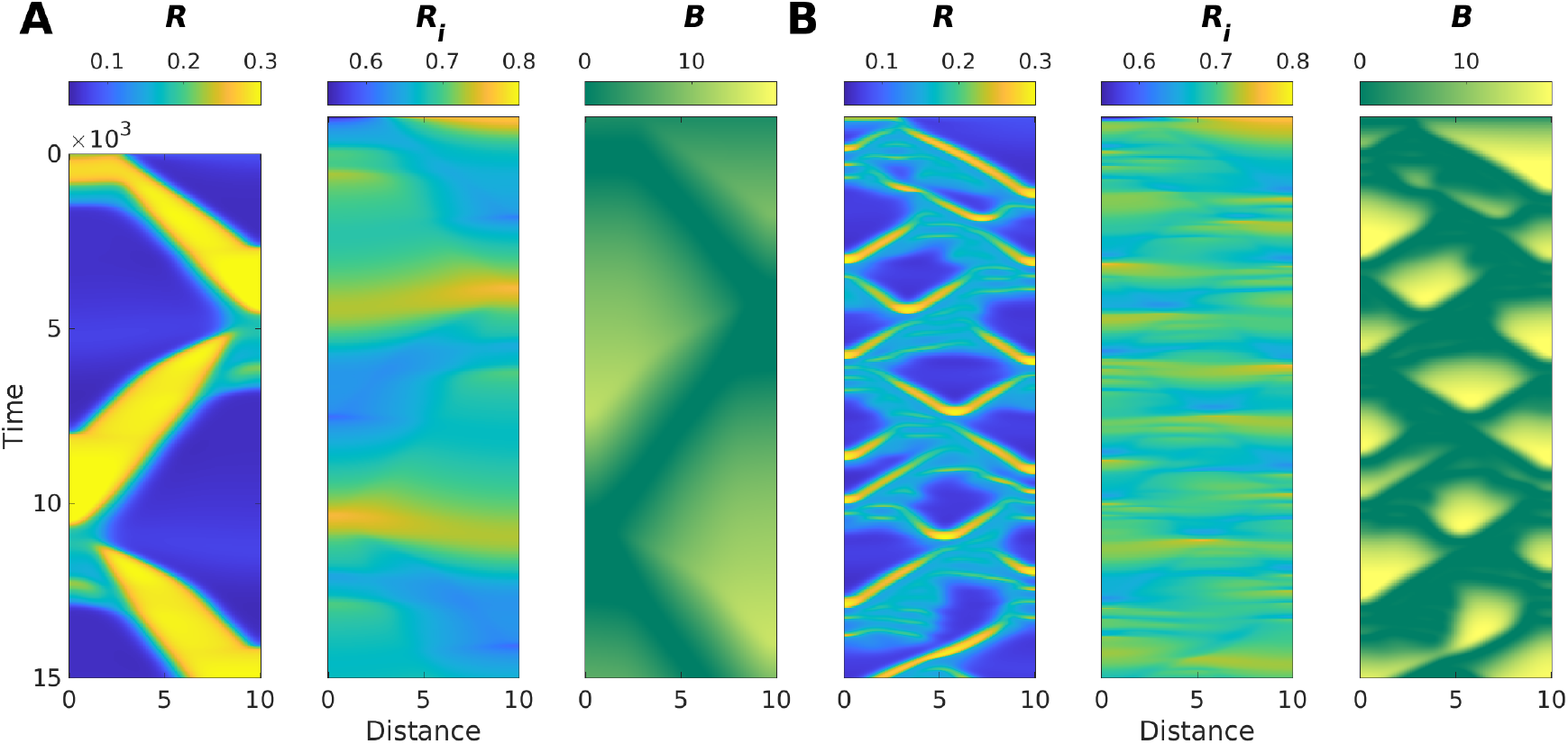
Switching and ripples produced by the 4D PDE model, given by 15, when *k_B_* is set to a constant. Kymographs of *R* (left) *R_i_* (middle) and *B* (right) in two parameter regimes defined by *k_B_*: (A) The RO regime and (B) SAO regime. A step-function is used as an initial condition, with zero-flux boundary conditions. The kymographs are color-coded according to the color-map in each panel.

The fact that *k_B_* varies slowly between these two dynamic behaviours in the 4D PDE model, given by Eqs. (15), indicates that a mixture of these outcomes should be observed periodically. In other words, we expect the 4D PDE model to show a succession of the behaviours detected in the two regimes displayed in Figure 11. Such a mixture of dynamic outcomes can potentially act as a framework that allows cells to exhibit not only change in Rac localization, but also membrane oscillations and fast directionality variations.

## 5 Computer simulations of motile cells

Cellular Potts Models (CPMs) have recently received quite a bit of attention as a tool to simulate “cells” in two-dimensional grid to investigate how different mechanical and chemical forces affect cellular migration [30,43,44]. It is an iterative discrete grid-based stochastic simulation technique that involves the modelling of the extracellular matrix (ECM) as a mesh upon which simulated cells are superimposed. This computational technique is used here as a framework to explore the migration patterns that can be generated by the 4D PDE model in combination with stochasticity.

### 5.1 Numerical implementation of the CPM

Each simulated CPM cell is modelled as a compartmentalized object that can grow, undergo shape variations and move on the surface of the grid. To account for forces **F** affecting cell shape and movement, such as elastic restoring forces or pressure caused by actin filaments growth that push the membrane outward, a Hamiltonian function *H*(**F**; ***λ***) is designed with different weights given by ***λ***. Monte Carlo Markov Chain-like simulations are then implemented based on the energy expressed by the Hamiltonian, to generate successive configurations of the cell body. Such configurations dictate how cell membrane evolves over time according to the physical forces included in the Hamiltonian. One can use *Morpheus*, a highly flexible CPM simulator software [45], to incorporate multi-scale systems into this computational technique to simulate a migrating cell.

In the analysis presented here, we combine the 4D PDE model, given by Eqs. (15), with the CPM simulations. This is done by incorporating the chemical reactions of the PDE model along with diffusion to dictate how membrane protrusions are regulated. The main assumption made in these simulations is that elevated Rac causes localized membrane protrusions. In our settings, we set the Hamiltonian function to be

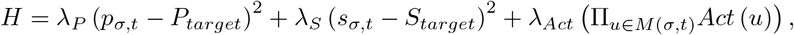

where *σ* denotes the configuration of the cell at time *t*, the weights *λ_P_* and *λ_S_* describe the deformation resistance with values tuned to allow the CPM cell to exhibit reasonable membrane elasticity properties [46, 47], *M* (*σ, t*) denotes the set of lattices that form the membrane of the cell and the *Act* function measures the amount of actin activity at a specific location. According to this model, protrusions are favoured in the regions of elevated Rac thanks to the *Act* function. Details of the Morpheus implementation can be found in [45].

### 5.2 Outcomes of the CPM simulations

As highlighted previously, the 4D PDE model, given by Eqs. 15, possesses different time scales; the presence of these time scales have been shown to be crucial for producing complex spatiotemporal patterns, induced by a combination of ROs and SAOs in the form of MMOs. It would be thus interesting to explore how these patterns affect migration paths of simulated cells generated using the CPM.

To study the migration paths of CPM cells, the boundary of these cells is initially computed using binary masks, defined as the black and white pixelation of the domain where each pixel occupied (unoccupied) by the cell is colored black (white), and their center of mass is determined at each frame generated by the Monte Carlo simulations. The *x* and *y* position of the center of mass of each frame is then tracked to generate the migration path. With this analysis, we have found that these paths display three distinct migration patterns. In the first one, CPM cells exhibit the classic *directional* motion that is typically associated with wave-pinning (Fig. 12A). These cells are fast, unidirectional (i.e., rush in one direction) and possess a stable front (back) with high (low) *R*. In the second pattern, simulated CPM cells display a homogeneous cytoplasm in terms of *R* and are non-motile (Fig. 12B). They nonetheless exhibit fluctuations in their membrane and small divergence away from their initial position due to the inherent stochasticity of the Monte Carlo simulation. Finally, in the third pattern, simulated cells show oscillations as well as meandering motion (Fig. 12C). More specifically, these cells protrude episodically and move more substantially than those simulated cells detected in the second pattern. Their membranes display waving patterns, and sometimes stretch because of multiple protruding fronts that dynamically change localization. No straight migratory patterns emerge in these CPM cells, because of this episodic activity in protrusions and retractions. This causes the simulated cells in this migration pattern to exhibit meandering trajectories typical of many migrating mesenchymal cells (such as CHO-K1 cells) [31]. This seems to suggest that MMOs are important for generating such migration pattern.

**Figure 12:**
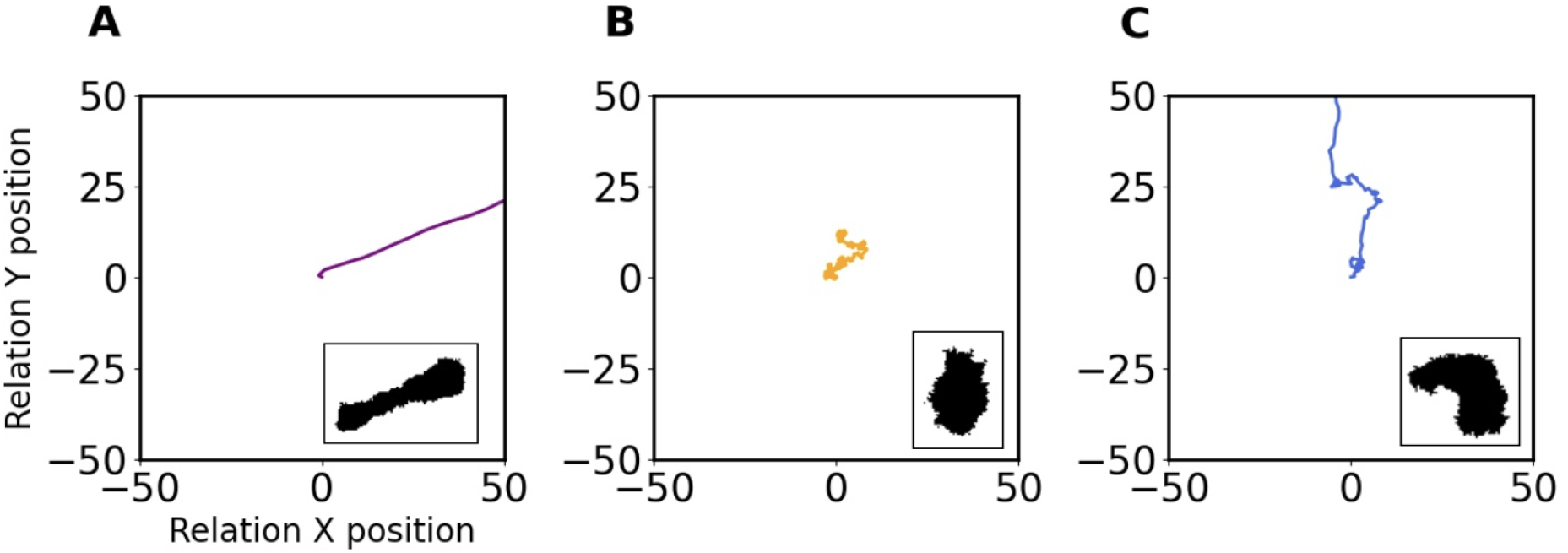
Migration patterns detected in the CPM. Migration paths of simulated CPM cells exhibiting (A) directed, (B) non-motile and (C) meandering motion. Insets: Binary masks of CPM cells at a given frame of the Monte Carlo simulations.

## 6 Discussion

Understanding molecular mechanisms underlying excitability in cell motility is a very challenging task due to the wide variability and complexity of the biological network involved. Redundancy in the signalling pathways, stochasticity in molecular expression and the nonlinearity associated with the processes underlying cell motility make investigating the role of excitability in this system very challenging experimentally. Therefore, one can use mathematical modeling as an alternative approach to decipher the dynamics of cell motility, including excitability, in different conditions.

In this present study, we used this modeling approach to uncover the oscillatory mechanisms that can arise in a molecularly-explicit model of cell motility. By focusing on a pre-existing 6D PDE model that describes closely the dynamics of the scaled concentrations of active and inactive forms of Rac and Rho (two members of the Rho family of GTPases that define cellular polarity), as well as of phosphorylated and unphosphorylated forms of the adaptor protein paxillin (an adhesion protein) [26], we were able to initially simply the model into a 1D ODE model by ignoring diffusion and by using a combination of linear regression and QSSA. The 1D ODE model described the dynamics of scaled active Rac concentration (*R*) and produced outcomes very similar to those produced by the original PDE model when diffusion was ignored, including bistability.

The simplified model was then successively expanded by phenomenologically turning two of its key parameters to slow and very slow variables. This expansion allowed the resulting 2D and 3D ODE models to produce ROs and MMOs in addition to bistability seen in the original model. The two parameters targeted in these expansions were the phosphorylation rate of paxillin *B* and the recovery rate *k_B_*. By applying slow fast analysis on the 2D and 3D ODE model, we determined how these oscillations are regulated by the critical manifold of this system and by the different time scales included. More specifically, by focusing on the 3D ODE model, we demonstrated how SAOs and LAOs in MMOs seen in scaled active Rac concentration were generated by considering the layer and reduced problems obtained from the timescale separation. Based on this analysis, we concluded that MMOs are due to both canard explosion and delayed Hopf bifurcation [32].

Whereas MMOs usually arise from folded singularities [32], the 3D ODE model analyzed here behaved differently. By choosing to decipher the dynamics of this model using two time scales: two fast (*R* and *B*) and one slow (*k_B_*) variables, we showed that the SAOs in MMOs start with very low amplitude but gradually grow before entering the RO regime; during this process the oscillations evolve from being canards without head to becoming canards with head in the RO regime in a manner similar to that governed by delay in a dynamical Hopf bifurcation [32]. By expanding the analysis to two time scales: one fast (*R*) and one slow (*B*), we then demonstrated that the last LAO in the MMOs was due to slow propagation of the trajectory along the two attracting sheets of the critical manifold and jump between them across the *B* nullcline.

While it is generally accepted that the small GTPases Rac and Rho are responsible for cell motility, most of the standard existing mathematical models only account for polarized cells with a leading front and a stable back that do not display a behavior more complex than straight directed motion. Some relevant models happen to produce oscillations [29, 37], but they rely on external feedbacks such as adhesion dynamic and signalling from the extra-cellular matrix, as well as do not exhibit frequent directionality change. Here we choose to focus only on internal pathways, and take advantage of the unusual structure and shape of the critical manifold and of the time-scale separation to show that these three internal variables are sufficient to build a model capable of generating complex oscillatory phenomena. This was verified when simulating the 2D and 4D PDE models considered in this study. The spatiotemporal patterns generated by the latter especially revealed that a mixture of switching between front and back as well as ripples across the one-dimensional domain can be obtained. This allowed the simulated CPM cell to exhibit three migration patterns that are quite distinct: directed, meandering and non-motile motions. The latter two appeared quite consistent with those migration patterns detected in CHO-K1 cells [31].

What are the functional and physiological consequences of the MMOs in this chemical system of cell motility is still to be determined. One needs to quantitatively compare the migration patterns generated by the CPM cells with those generated by mesenchymal cells (e.g. CHO-K1 cells), in a rigourous manner that should be applicable to both systems. This would allowfor important conclusions to be drawn about the role of MMOs in defining cell migration.

## References

[1] Eugene M Izhikevich. Neural excitability, spiking and bursting. International journal of bifurcation and chaos, 10(06):1171–1266, 2000.

[2] John Rinzel and G Bard Ermentrout. Analysis of neural excitability and oscillations. Methods in neuronal modeling, 2:251–292, 1998.

[3] Rob J De Boer and Alan S Perelson. Target cell limited and immune control models of hiv infection: a comparison. Journal of theoretical Biology, 190(3):201–214, 1998.

[4] Steven A Prescott. Excitability: Types i, ii, and iii., 2014.

[5] Steven A Prescott, Yves De Koninck, and Terrence J Sejnowski. Biophysical basis for three distinct dynamical mechanisms of action potential initiation. PLoS computational biology, 4(10):e1000198, 2008.

[6] Zhiguo Zhao, Li Li, and Huaguang Gu. Different dynamical behaviors induced by slow excitatory feedback for type ii and iii excitabilities. Scientific Reports, 10(1):1–16, 2020.

[7] Alan L Hodgkin. The local electric changes associated with repetitive action in a non-medullated axon. The Journal of physiology, 107(2):165, 1948.

[8] Ilan D Zipkin, Rachel M Kindt, and Cynthia J Kenyon. Role of a new rho family member in cell migration and axon guidance in c. elegans. Cell, 90(5):883–894, 1997.

[9] Elena Scarpa and Roberto Mayor. Collective cell migration in development. Journal of Cell Biology, 212(2):143–155, 2016.

[10] Li Li, Yong He, Min Zhao, and Jianxin Jiang. Collective cell migration: Implications for wound healing and cancer invasion. Burns & trauma, 1(1):2321–3868, 2013.

[11] Douglas Hanahan and Robert A Weinberg. Hallmarks of cancer: the next generation. cell, 144(5):646–674, 2011.

[12] Atef Asnacios and Olivier Hamant. The mechanics behind cell polarity. Trends in cell biology, 22(11):584–591, 2012.

[13] Athanasius FM Marée, Alexandra Jilkine, Adriana Dawes, Verônica A Grieneisen, and Leah Edelstein-Keshet. Polarization and movement of keratocytes: a multiscale modelling approach. Bulletin of mathematical biology, 68(5):1169–1211, 2006.

[14] Erin L Barnhart, Jun Allard, Sunny S Lou, Julie A Theriot, and Alex Mogilner. Adhesion-dependent wave generation in crawling cells. Current Biology, 27(1):27–38, 2017.

[15] Laurent MacKay, Etienne Lehman, and Anmar Khadra. Deciphering the dynamics of lamellipodium in a fish keratocytes model. Journal of Theoretical Biology, 512:110534, 2021.

[16] Sabina E Winograd-Katz, Reinhard Fässler, Benjamin Geiger, and Kyle R Legate. The integrin adhesome: from genes and proteins to human disease. Nature reviews Molecular cell biology, 15(4):273–288, 2014.

[17] Satyajit K Mitra, Daniel A Hanson, and David D Schlaepfer. Focal adhesion kinase: in command and control of cell motility. Nature reviews Molecular cell biology, 6(1):56–68, 2005.

[18] Cord Brakebusch and Reinhard Fässler. The integrin–actin connection, an eternal love affair. The EMBO journal, 22(10):2324–2333, 2003.

[19] Catherine D Nobes and Alan Hall. Rho, rac, and cdc42 gtpases regulate the assembly of multimolecular focal complexes associated with actin stress fibers, lamellipodia, and filopodia. Cell, 81(1):53–62, 1995.

[20] Anne J Ridley, Hugh F Paterson, Caroline L Johnston, Dagmar Diekmann, and Alan Hall. The small gtp-binding protein rac regulates growth factor-induced membrane ruffling. Cell, 70(3):401–410, 1992.

[21] Anne J Ridley. Rho gtpases and cell migration. Journal of cell science, 114(15):2713–2722, 2001.

[22] Sarah J Heasman and Anne J Ridley. Mammalian rho gtpases: new insights into their functions from in vivo studies. Nature reviews Molecular cell biology, 9(9):690–701, 2008.

[23] Anjana Nayal, Donna J Webb, Claire M Brown, Erik M Schaefer, Miguel Vicente-Manzanares, and Alan Rick Horwitz. Paxillin phosphorylation at ser273 localizes a git1–pix–pak complex and regulates adhesion and protrusion dynamics. The Journal of cell biology, 173(4):587–589, 2006.

[24] Keith Burridge and Krister Wennerberg. Rho and rac take center stage. Cell, 116(2):167–179, 2004.

[25] Abira Rajah, Colton G Boudreau, Alina Ilie, Tse-Luen Wee, Kaixi Tang, Aleksandar Z Borisov, John Orlowski, and Claire M Brown. Paxillin s273 phosphorylation regulates adhesion dynamics and cell migration through a common protein complex with pak1 and *β*pix. Scientific reports, 9(1):1–20, 2019.

[26] Kaixi Tang, Colton G Boudreau, Claire M Brown, and Anmar Khadra. Paxillin phosphorylation at serine 273 and its effects on rac, rho and adhesion dynamics. PLoS computational biology, 14(7):e1006303, 2018.

[27] William R Holmes, May Anne Mata, and Leah Edelstein-Keshet. Local perturbation analysis: a computational tool for biophysical reaction-diffusion models. Biophysical journal, 108(2):230–236, 2015.

[28] Sulagna Das, Taofei Yin, Qingfen Yang, Jingqiao Zhang, Yi I Wu, and Ji Yu. Single-molecule tracking of small gtpase rac1 uncovers spatial regulation of membrane translocation and mechanism for polarized signaling. Proceedings of the National Academy of Sciences, 112(3):E267–E276, 2015.

[29] Elisabeth G Rens and Leah Edelstein-Keshet. Cellular tango: how extracellular matrix adhesion choreographs rac-rho signaling and cell movement. Physical biology, 18(6):066005, 2021.

[30] Athanasius FM Marée, Verônica A Grieneisen, and Paulien Hogeweg. The cellular potts model and biophysical properties of cells, tissues and morphogenesis. In Single-cell-based models in biology and medicine, pages 107–136. Springer, 2007.

[31] Lucie Plazen, Jalal Al Rahbani, Claire M Brown, and Anmar Khadra. Polarity and mixed-mode oscillations may underlie different patterns of cellular migration. Scientific Report, under review, 2022.

[32] Mathieu Desroches, John Guckenheimer, Bernd Krauskopf, Christian Kuehn, Hinke M Osinga, and Martin Wechselberger. Mixed-mode oscillations with multiple time scales. Siam Review, 54(2):211–288, 2012.

[33] Mathieu Desroches, Tasso J Kaper, and Martin Krupa. Mixed-mode bursting oscillations: Dynamics created by a slow passage through spike-adding canard explosion in a square-wave burster. Chaos: An Interdisciplinary Journal of Nonlinear Science, 23(4):046106, 2013.

[34] Peter Szmolyan and Martin Wechselberger. Canards in r3. Journal of Differential Equations, 177(2):419–453, 2001.

[35] Valery Petrov, Stephen K Scott, and Kenneth Showalter. Mixed-mode oscillations in chemical systems. The Journal of chemical physics, 97(9):6191–6198, 1992.

[36] Jessica K Lyda, Zhang L Tan, Abira Rajah, Asheesh Momi, Laurent Mackay, Claire M Brown, and Anmar Khadra. Rac activation is key to cell motility and directionality: an experimental and modelling investigation. Computational and Structural Biotechnology Journal, 17:1436–1452, 2019.

[37] William R Holmes, JinSeok Park, Andre Levchenko, and Leah Edelstein-Keshet. A mathematical model coupling polarity signaling to cell adhesion explains diverse cell migration patterns. PLoS computational biology, 13(5):e1005524, 2017.

[38] Yoichiro Mori, Alexandra Jilkine, and Leah Edelstein-Keshet. Wave-pinning and cell polarity from a bistable reaction-diffusion system. Biophysical journal, 94(9):3684–3697, 2008.

[39] Eusebius J Doedel, Volodymyr A Romanov, Randy C Paffenroth, Herbert B Keller, Donald J Dichmann, Jorge Galán-Vioque, and André Vanderbauwhede. Elemental periodic orbits associated with the libration points in the circular restricted 3-body problem. International Journal of Bifurcation and Chaos, 17(08):2625–2677, 2007.

[40] R Bertram et al. Topological and phenomenological classification of bursting oscillations. Bulletin of mathematical biology, 57(3):413–39, 1995. Mathematical Neuroscience.

[41] David G Drubin and W James Nelson. Origins of cell polarity. Cell, 84(3):335–344, 1996.

[42] Yoichiro Mori, Alexandra Jilkine, and Leah Edelstein-Keshet. Asymptotic and bifurcation analysis of wave-pinning in a reaction-diffusion model for cell polarization. SIAM Journal on applied mathematics, 71(4):1401–1427, 2011.

[43] Elisabeth G Rens and Leah Edelstein-Keshet. From energy to cellular forces in the cellular potts model: An algorithmic approach. PLoS computational biology, 15(12):e1007459, 2019.

[44] Noriyuki Bob Ouchi, James A Glazier, Jean-Paul Rieu, Arpita Upadhyaya, and Yasuji Sawada. Improving the realism of the cellular potts model in simulations of biological cells. Physica A: Statistical Mechanics and its Applications, 329(3-4):451–458, 2003.

[45] Jörn Starruß, Walter de Back, Lutz Brusch, and Andreas Deutsch. Morpheus: a user-friendly modeling environment for multiscale and multicellular systems biology. Bioinformatics, 30(9):1331–1332, 01 2014.

[46] François Graner and James A. Glazier. Simulation of biological cell sorting using a two-dimensional extended potts model. Physical Review Letters, 69(13):2013–2016, 1992.

[47] Ioana Niculescu, Johannes Textor, and Rob J. de Boer. Crawling and gliding: A computational model for shape-driven cell migration. PLOS Computational Biology, 11, 10 2015.

[48] Clare M Waterman-Storer and ED Salmon. Positive feedback interactions between microtubule and actin dynamics during cell motility. Current opinion in cell biology, 11(1):61–67, 1999.

[49] Martin Krupa and Martin Wechselberger. Local analysis near a folded saddle-node singularity. Journal of Differential Equations, 248(12):2841–2888, 2010.

[50] Rodica Curtu. Singular hopf bifurcations and mixed-mode oscillations in a two-cell inhibitory neural network. Physica D: Nonlinear Phenomena, 239(9):504–514, 2010.

[51] Leah Edelstein-Keshet, William R Holmes, Mark Zajac, and Meghan Dutot. From simple to detailed models for cell polarization. Philosophical Transactions of the Royal Society B: Biological Sciences, 368(1629):20130003, 2013.

[52] Alan Mathison Turing. The chemical basis of morphogenesis. Bulletin of mathematical biology, 52(1):153–197, 1990.

[53] Bard Ermentrout and Ajay Mahajan. Simulating, analyzing, and animating dynamical systems: a guide to xppaut for researchers and students. Appl. Mech. Rev., 56(4):B53–B53, 2003.

